# Deep Learning with Evolutionary Scale Modeling to discover Antibiotic-Resistance Drug against Klebsiella Pneumoniae

**DOI:** 10.1101/2024.11.09.622818

**Authors:** Linxiao Jin, Zhihao Lei, Peng Lin, Cong Ma, Wen Zhang, Jiali Zhou, Sijun Meng, Zhenghua Jiang, Yemei Bu, Yingfan Xu, Nijun Wei, Tsehao Hsu, Glen Qin, Hesong Qiu

## Abstract

The development of effective antibiotics is crucial in combating antibiotic-resistant pathogens such as Klebsiella pneumoniae (KP). This study utilizes deep learning, specifically Evolutionary Scale Modeling (ESM), to identify potential drug candidates. By analyzing molecular similarities between known drugs and KP strains, we aim to accelerate the discovery of new treatments and reduce the risks associated with clinical trials. This approach may shed new lights in the field of drug repositioning.

## 1. Introduction

The emergence and rapid spread of antibiotic-resistant pathogens represent one of the greatest challenges in modern healthcare, with Klebsiella pneumoniae (KP) standing out due to its high resistance to multiple antibiotics. This pathogen is particularly problematic in hospital settings, where it causes severe infections with limited treatment options. Current antibiotic treatments for Klebsiella pneumoniae face several significant challenges and limitations.

One of the primary issues is the high level of antibiotic resistance exhibited by Klebsiella pneumoniae. Multidrug-resistant (MDR) strains exhibit resistance to several classes of antibiotics, including β-lactams, aminoglycosides, and quinolones, making standard therapies ineffective ^[1]^. Particular concern is rising resistance to carbapenems and colistin, which are two critical last-resort antibiotics. Resistance to carbapenems has reached alarming levels with rates as high as 95% in some regions, which is largely due to carbapenemase-producing strains like Klebsiella pneumoniae carbapenemase (KPC) ^[2]^. Similarly, colistin resistance has surged dramatically, and some studies report an increase from 8% to 72% over a few years ^[3]^.

In response to this resistance, combination therapies involving multiple antibiotics are often employed. However, these combinations frequently yield suboptimal results ^[1]^. Common regimens include high-dose meropenem, colistin, fosfomycin, tigecycline, and aminoglycosides. Despite their widespread usage, these therapies might be inconsistent in effectiveness, which may not always prevent the emergence of resistance. Additionally, the antibiotics used in these combinations can cause significant toxicity, including nephrotoxicity and neurotoxicity, which limits their applications in clinics ^[2]^.

The slow pace of new antibiotic development further complicates treatment strategies for KP. The development pipeline for new antibiotics has not kept up with the rapid emergence of resistant strains. Although some new antimicrobials targeting MDR-KP are under development, they are still in various stages of clinical research, which are not yet widely available ^[6]^. Moreover, even newly developed antibiotics can quickly encounter resistance, as seen with drugs like tigecycline and minocycline, which were initially effective against MDR strains but have faced resistance issues ^[3]^.

KP’s genetic diversity and its ability to acquire resistance genes through horizontal gene transfer exacerbate the problem. The bacterium possesses a wide array of resistance genes for β-lactamases, carbapenemases, and efflux pumps, contributing to its formidable resistance profile ^[4]^. Its capability to acquire resistance genes from other bacteria through horizontal gene transfer further spreads resistance mechanisms, making it a formidable opponent in the fight against antibiotic resistance.

Preventing the spread of MDR and extensively drug-resistant (XDR) KP in healthcare settings requires rigorous infection control measures, which can be challenging to implement consistently. Effective antimicrobial stewardship programs are essential to limit the use of broad-spectrum antibiotics, which reduce the selection pressure for resistant strains. Nevertheless, the implementations of these programs face challenges due to varying levels of resources and adherence to guidelines ^[5]^.

The global prevalence of antibiotic-resistant KP varies significantly across different regions, with a notable increase in MDR and XDR strains globally. In China, for instance, the isolation rate of KP surged from 1.57% in 2006 to 32.30% in 2020, reflecting broader regional issues ^[9]^. This rise necessitates urgent surveillance and control measures across Asia. A systematic review and meta-analysis estimated the global prevalence of nosocomial MDR KP at 32.8%, underscoring its significant presence in healthcare settings worldwide ^[8]^. The rise of antibiotic-resistant KP has profound implications for public health, influencing both clinical outcomes and healthcare systems. Infections caused by MDR and XDR strains of KP are associated with high mortality rates, sometimes exceeding 40% within 30 days of infection. These infections typically occur amongst vulnerable populations, such as those in intensive care units (ICUs) or with underlying health conditions ^[7]^. Patients infected with resistant strains often require longer hospital stays, increasing the risk of further complications and nosocomial infections.

The economic burden of treating infections caused by resistant KP strains is significant. Treatment is more expensive due to the need for potent, often toxic antibiotics, extended hospital stays, and additional infection control measures. For example, in Brazil ^[7]^, the direct drug cost per patient infected by KPC is estimated to be nearly $4,100. The economic burden strains healthcare resources, diverting funds from other essential services and potentially compromising the overall quality of care.

Robust surveillance systems are crucial to monitor the spread of resistant KP strains and implement timely interventions. Initiatives like the Global Antimicrobial Resistance and Surveillance System (GLASS) emphasize the importance of such measures ^[11]^. Strengthening infection control practices in healthcare settings, such like rigorous hygiene protocols and isolation measures, is essential to prevent the spread of resistant strains.

Implementing antimicrobial stewardship programs to promote the judicious use of antibiotics can help reduce the selection pressure that drives the emergence of resistant strains. This includes guidelines for appropriate antibiotic prescribing and de-escalation of therapy based upon cultures ^[10]^. Additionally, there is an urgent demand for the development of new antibiotics and alternative therapies to combat resistant KP strains. Investment in research and development is crucial to stay ahead of evolving resistance mechanisms.

The global prevalence of antibiotic-resistant KP and its impact on public health are significant, which attracts growing concerns. Addressing these challenges requires a multifaceted approach, including enhanced surveillance, stringent infection control practices, rational antibiotic use, and the development of new therapeutic options. The collective efforts of healthcare providers, policymakers, and researchers are crucial to mitigate the threat posed by this formidable pathogen.

## 2 Materials and Methods

### 2.1 Datasets

Two primary datasets were utilized for this investigation:

#### a) DrugBank Database ^[17]^

This comprehensive resource provides detailed information about drugs and their targets, including data on structures, enzymes, and pathways. For this study, the DrugBank database was meticulously filtered to focus on biotech and small molecule drugs, specifically excluding vaccines, antibodies, toxins, and other irrelevant categories. In addition to these exclusions, we also eliminated drugs known to easily interact with others, as outlined in Table 1. This exclusion was crucial because we aimed to avoid selecting drugs that could potentially cause adverse reactions when used in combination, a key consideration for the deep learning model. After this thorough filtering process, we refined the dataset to include 3,475 unique drugs, from which key features such as targets, enzymes, SMILES representations, and pathways were extracted for subsequent analysis.

**Table 1.**
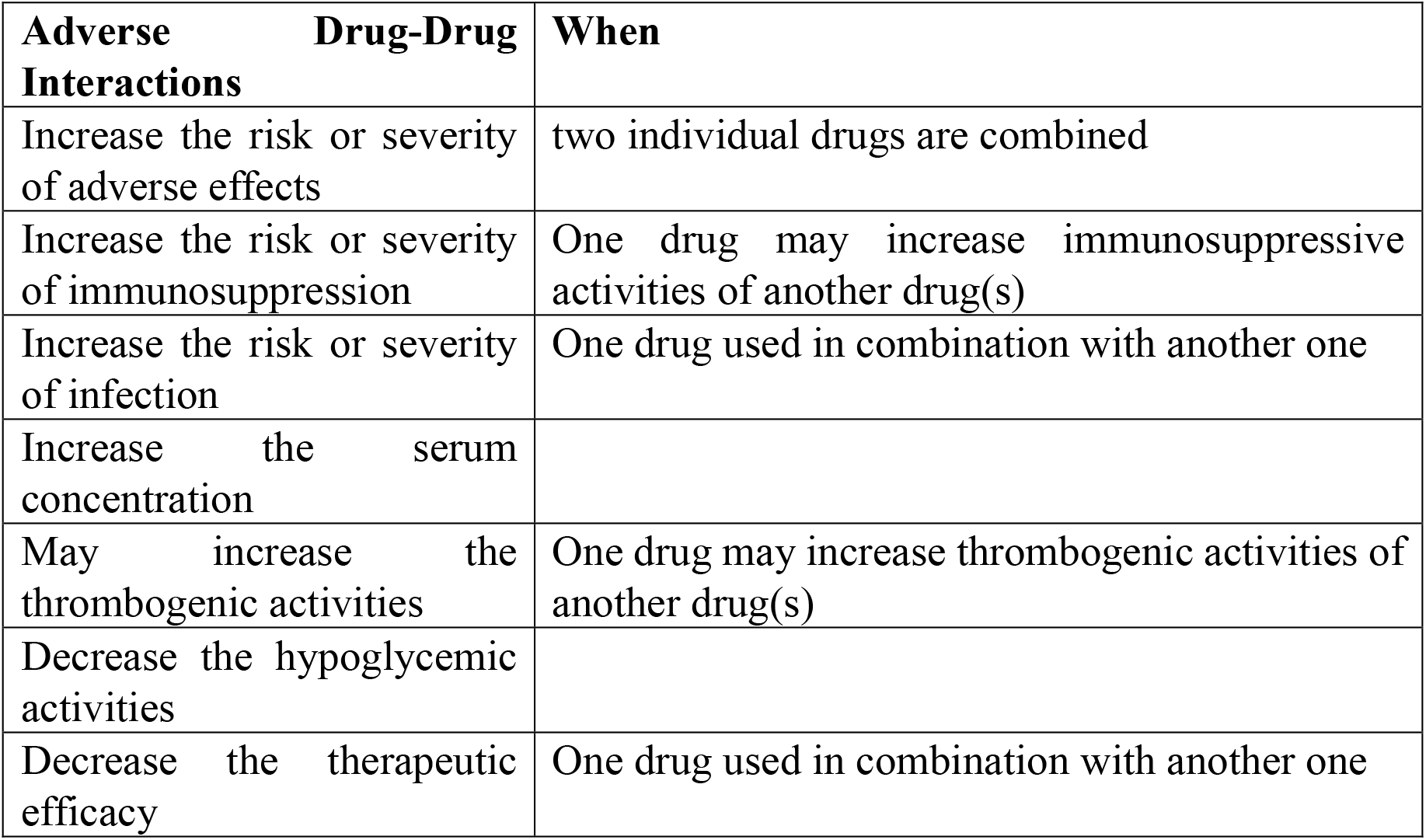
Adverse drug-drug interactions when 2 drugs combined or utilized.

#### b) KP Strain Fasta Dataset ^[18]^

It consists of 831 KP strains collected from patients in hospitals, which is used to extract target proteins for similarity analysis. From 831 KP strains we have extracted about 220,000 protein targets.

### 2.2 Molecular Similarity Comparison

In this study, we leveraged the power of Evolutionary Scale Modeling (ESM) ^[13]^ to conduct a comprehensive molecular similarity comparison, focusing on the structural analysis of drugs and Klebsiella pneumoniae (KP) target proteins. The ESM-2 model, along with ESMFold, represents a significant advancement in protein language models, offering unique capabilities for predicting protein structures from single-sequence inputs. This approach simplifies the prediction process and reduces computational requirements compared to models like AlphaFold2 and RoseTTAFold, which rely heavily on multiple sequence alignments (MSAs) and templates ^[12]^.

The key advantage of using ESMFold lies in its ability to generate accurate structure predictions directly from a single sequence, making it considerably faster than alignment-based methods. This speed advantage has facilitated the creation of extensive databases such as the ESM Metagenomic Atlas, containing over 600 million metagenomic protein structures. While other models like AlphaFold2 and RoseTTAFold achieve high accuracy, they are computationally intensive and slower due to their reliance on MSAs and template-based approaches ^[12]^. Ensemble models like ProtENN also perform well in specific tasks but may not match the overall speed and scalability of ESM-2.

To conduct an accurate and large-scale molecular similarity analysis, we employed the ESMFold model to generate normalized embedding vectors for both the drugs listed in the DrugBank database and the target proteins of 831 Klebsiella pneumoniae (KP) strains. These normalized vectors allowed for a direct and unbiased comparison through cosine similarity measures, facilitating a precise assessment of molecular resemblance. The efficient handling of large datasets was made possible through this approach, enabling the rapid processing of the data necessary for large-scale screening. Additionally, GPU acceleration significantly enhanced the speed of matrix calculations, allowing us to process approximately 484 million pairwise comparisons efficiently.

Given the vast scale of the dataset, we initially validated our algorithm using a smaller subset, which comprised one-tenth of the total strains. This preliminary testbed was crucial in refining our methodology and ensuring the algorithm’s effectiveness. In particular, we focused on identifying the top 0.01% of similarity pairs, a strategy that allowed us to filter out less relevant pairings. By implementing a minimal heap or priority queue, we were able to retain only the most significant similarity pairs, concentrating our efforts on the most promising drug-bacteria combinations. This targeted approach not only accelerated processing times but also enhanced our ability to identify potential drug candidates more effectively. Figure 1 illustrates the overall process flow of ESM, depicting the comparison between the 831 bacterial strains and the 3,475 drugs with their respective features.

**Figure 1.**
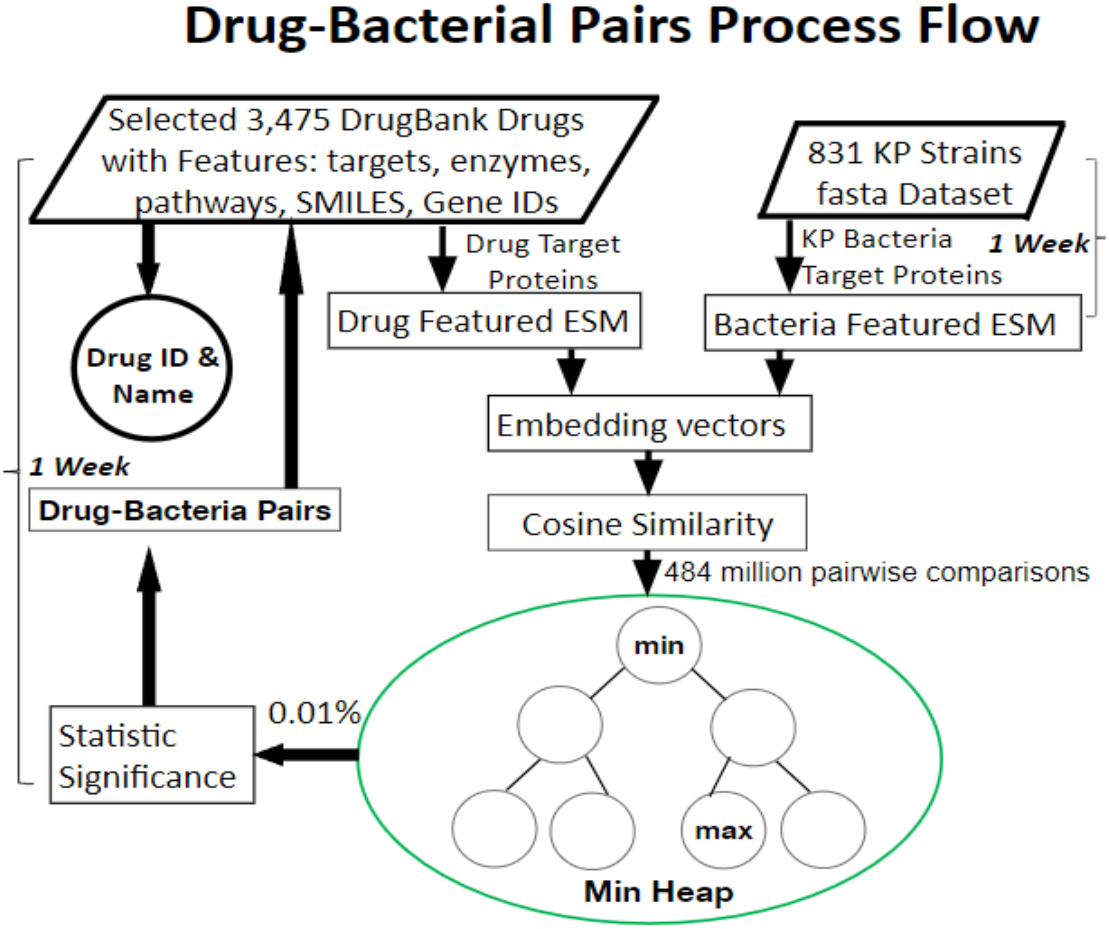
Process Flow of Drug-Bacterial Pairs Similarity Analysis.

To assess molecular similarity, we utilized cosine similarity measures due to their several advantages in this context. Cosine similarity measures the cosine of the angle between two vectors in a multi-dimensional space, focusing on the orientation rather than the magnitude of the vectors. This makes it particularly effective for comparing the shape and structure of molecular features without being influenced by their size. The simplicity and interpretability of cosine similarity make it straightforward to compute using vector representations of molecular fingerprints. Additionally, cosine similarity is well-suited for high-dimensional data, such as molecular fingerprints that capture various structural features of molecules.

The process involved generating embedding vectors for each drug and each KP target protein, followed by pairwise comparisons using cosine similarity. Each comparison measured the angle between the vectors, providing a reliable metric for gauging the degree of structural similarity. This method was particularly useful for filtering out less promising candidates and focusing on those with higher potential for therapeutic efficacy.

While alternative metrics for molecular similarity, such as the Tanimoto (Jaccard) Coefficient, Dice Coefficient ^[15]^, and Spec2Vec ^[14]^, each have their strengths, we chose cosine similarity for its straightforward implementation and efficiency. The computational efficiency of cosine similarity, especially when accelerated by GPUs, made it the optimal choice for our large-scale screening needs.

Overall, the use of ESM and cosine similarity in this study provided a robust framework for identifying potential drug candidates against Klebsiella pneumoniae. The efficiency and accuracy of these methods offer a significant advantage in the fast-paced field of drug discovery, particularly in the context of combating antibiotic resistance.

## 3 Results

### 3.1 Data Extraction

From the initial 3,475 unique drugs, a subset with relevant features (targets, enzymes, SMILES, and pathways) was identified. Results in Table 2 illustrated how many unique drugs were identified for each respective key feature. ESM modeling for similarity analysis narrows down potential drug candidates based solely on the 2,155 unique drugs with the Targets feature.

**Table 2.** Drug Count with respective Features among 3,475 Drugs.

| DrugBank<br>Drugs | Targets | Enzymes | SMILE<br>S | Pathways |
| --- | --- | --- | --- | --- |
| 3,475 | 2,155 | 889 | 3,017 | 435 |

### 3.2 Molecular Similarity Analysis

To assess molecular similarity, we utilized cosine similarity measures due to their several advantages in this context. Cosine similarity (1) measures the cosine of the angle between two vectors in a multi-dimensional space, focusing on the orientation rather than the magnitude of the vectors. This makes it particularly effective for comparing the shape and structure of molecular features without being influenced by their size. The simplicity and interpretability of cosine similarity make it straightforward to compute using vector representations of molecular fingerprints. Additionally, cosine similarity is well-suited for high-dimensional data, such as molecular fingerprints that capture various structural features of molecules.

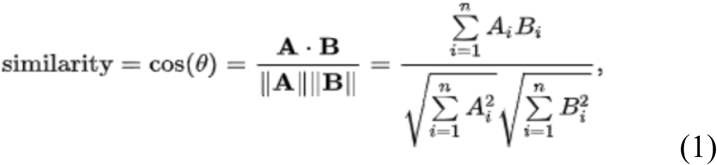

The ESMFold model was employed to generate normalized embedding vectors for both the drugs in the DrugBank database and the target proteins of 831 KP strains. These normalized vectors facilitated a direct and unbiased comparison through cosine similarity measures, allowing for an accurate assessment of molecular resemblance. This approach enabled us to handle large datasets efficiently and perform rapid comparisons necessary for large-scale screening. The use of GPU acceleration further sped up matrix calculations, enabling the processing of approximately 484 million pairwise comparisons in a timely manner.

The process involved generating embedding vectors for each drug and each KP target protein, followed by pairwise comparisons using cosine similarity. Each comparison measured the angle between the vectors, providing a reliable metric for gauging the degree of structural similarity. This method was particularly useful for filtering out less promising candidates and focusing on those with higher potential for therapeutic efficacy.

While alternative metrics for molecular similarity, such as the Tanimoto (Jaccard) Coefficient, Dice Coefficient, and Spec2Vec, each have their strengths, we chose cosine similarity for its straightforward implementation and efficiency. The computational efficiency of cosine similarity, especially when accelerated by GPUs, made it the optimal choice for our large-scale screening needs.

To gain deeper insights into the overall structural trends within the dataset, we recorded similarities for statistical distribution analysis. These distributions provided valuable information on the prevalence of specific similarity ranges across the entire dataset, which was visually represented in Figure 2. This visualization offered a clear perspective on the similarity trends across all drug-bacteria pairs, effectively streamlining the large dataset into a more comprehensible format. By highlighting the most prevalent similarity ranges, this approach facilitated a more focused analysis of common structural features.

**Figure 2.**
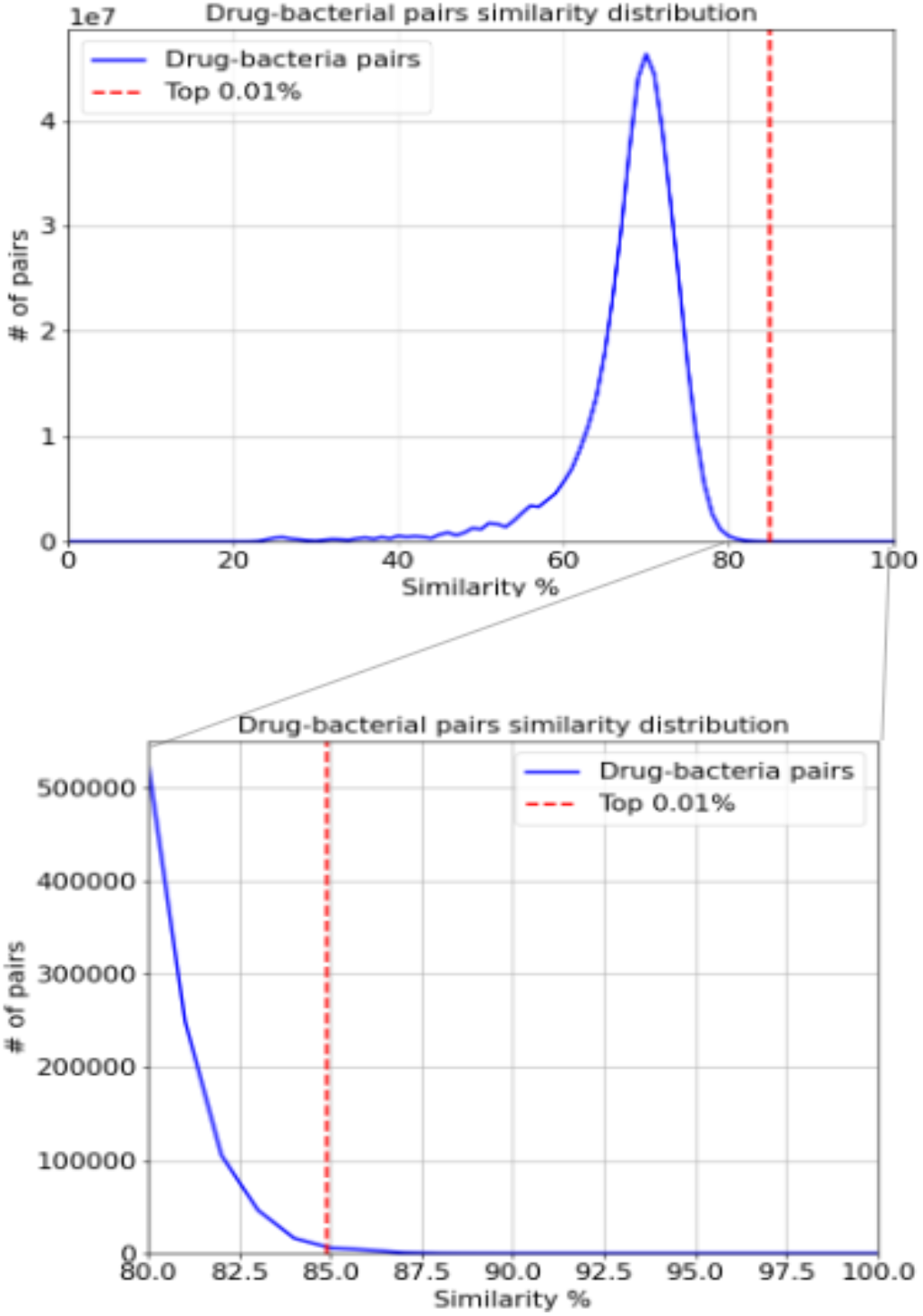
Drug-bacterial pairs distribution in similarity.

Given the substantial size of the dataset, we initially validated our algorithm on a smaller subset, consisting of one-tenth of the total strains, to ensure its effectiveness and efficiency. This smaller testbed allowed us to refine our methodology, particularly in identifying the top 0.01% of similarity pairs. By using a min-heap to focus on this top percentile, we were able to eliminate the long tail of less relevant pairs and concentrate on the most promising drug-bacteria pairings, thereby accelerating the processing time and enhancing the identification of potential drug candidates.

The results of the molecular similarity comparison are presented in Table 3. Notably, significant matches were identified with drugs such as Rofecoxib and Artenimol, which demonstrated high structural similarity to Klebsiella pneumoniae proteins. These findings are consistent with previous pharmacological studies that have recognized the efficacy of these drugs against Klebsiella pneumoniae. Additionally, other compounds, such as Radezolid, as well as elements like copper and zinc, appeared in high-similarity pairings, albeit less frequently. These observations are consistent with established therapeutic effects, indicating promising opportunities for leveraging these structural similarities in future drug design initiatives.

**Table 3.** Molecular Similarity Comparison Result.

| % of Top Similarity Pairs | Drug Target ID | DrugBank ID | Drug name |
| --- | --- | --- | --- |
| 47.47% | P15502 | DB00533 | Rofecoxib |
| 36.19% | Q8I6U8 | DB11638 | Artenimol |
| 6.59% | Q9P2E9 | DB12339 | Radezolid |
| 1.89% | P35527 | DB01593 & DB09130 | Zinc & Copper |
| 1.56% | Q8I487 | DB11638 | Artenimol |
| 6.3% | P51843<br>P10275<br>O43519<br>and other 21 target IDs | DB01234<br>DB00255<br>DB00619<br>and other 54 DrugBank IDs | Dexamethasone<br>Diethylstilbestrol<br>Imatinib<br>and other 54 drugs |

The high similarity scores between known effective drugs and Klebsiella pneumoniae (KP) proteins further validate the robustness of our ESM-based approach. Findings in other studies like ^[19]^, supports the validity of our predictions. This methodology not only identifies promising candidates for further testing but also offers a targeted strategy for drug repurposing. The strong correlation observed between structural similarity and therapeutic potential underscores the critical role of molecular modeling in accelerating drug discovery processes. As a next step, we are organizing clinical trials to confirm the efficacy of these predicted drugs in treating KP infections.

## 4 Conclusion and discussion

K. pneumoniae has been extensively studied for its diverse antibiotic resistance mechanisms, including the production of extended-spectrum β-lactamases (ESBLs), carbapenemases, and the presence of efflux pumps and porin mutations.

Whole genome sequencing (WGS) and genomic surveillance have been crucial in identifying high-risk clones and resistance genes. Similar genomic approaches can be applied to other antibiotic-resistant pathogens like Escherichia coli, Staphylococcus aureus, and Acinetobacter baumannii. By using ESM models to predict protein structures and identify resistance mechanisms, researchers can gain insights into how these pathogens evade antibiotics. For instance, WGS can help track the spread of resistance genes in E. coli and identify potential targets for new therapeutic interventions.

The findings underscore the importance of integrating advanced computational techniques in drug discovery. The use of ESM for molecular similarity analysis offers a robust framework for identifying effective treatments against resistant pathogens. Continued research in this area will further enhance our ability to combat antibiotic resistance and improve public health outcomes.

The ESM model has shown promise in predicting protein structures and aiding drug discovery. However, ESM models have limitations in accurately modeling certain protein structures, especially those with highly dynamic or disordered regions. These regions can be critical for drug binding and function, and inaccuracies in their modeling can lead to suboptimal drug design. Additionally, ESM models have billions of parameters, require significant computational resources for training and inference. This can be a barrier for smaller research institutions or companies with limited access to high-performance computing infrastructure.

This study demonstrates the potential of deep learning and ESM in identifying new drug candidates for treating antibiotic resistant KP infections. By focusing on molecular similarities, we can streamline the drug discovery process, significantly reducing the time and cost associated with traditional methods. Future work will involve rigorous preclinical and clinical trials to validate the efficacy of the identified drug candidates.

## Acknowledgements

This work was funded by Grant from Hong Kong Innovation and Technology Commission (grant no. PRP/062/22FX), Innovation Fund from Jinying Technology Co., Ltd. (grant no. CX: 20210506D) and Grant from Hong Kong Innovation and Technology Commission (grant no. PsH/072/22).

